# *In vivo* Monitoring of Glucose using Ultrasound-induced Resonance in Implantable Smart Hydrogel Microstructures

**DOI:** 10.1101/2021.04.11.439366

**Authors:** Navid Farhoudi, Lars B. Laurentius, Jules J. Magda, Christopher F. Reiche, Florian Solzbacher

**Affiliations:** Department of Electrical and Computer Engineering, University of Utah, Salt Lake City, 84112, USA; Department of Chemical Engineering, University of Utah, Salt Lake City, 84112, USA; Department of Materials Science & Engineering, University of Utah, Salt Lake City, UT 84112, USA; Department of Bioengineering, University of Utah, Salt Lake City, UT 84112, USA

**Keywords:** smart hydrogels, glucose-sensitive hydrogels, ultrasound imaging, implantable biomedical sensors, mechanical resonance, continuous glucose monitoring

## Abstract

A novel glucose sensor is presented that uses smart hydrogels as a biocompatible implantable sensing element, which completely eliminates the need for any implanted electronics and uses an external conventional medical-grade ultrasound transducer for readout. The readout mechanism makes use of resonance absorption of ultrasound waves in glucose-sensitive hydrogels. Changes in *in vivo* glucose concentration in the interstitial tissue lead to swelling and de-swelling of the gels which in turn lead to changes in resonance behavior. The hydrogels are designed and shaped such as to exhibit specific mechanical resonance frequencies while remaining sonolucent to other frequencies. Thus, they allow conventional and continued ultrasound imaging, while yielding a sensing signal at specific frequencies that is correlated with glucose concentration. The resonance frequencies can be tuned by changing the shape and mechanical properties of the gel structures, such as to allow for multiple, co-located implanted hydrogels with different sensing characteristics or targets to be employed and read out, without interference, using the same ultrasound transducer, by simply toggling frequencies. The fact that there is no need for any implantable electronics, also opens the path towards future use of biodegradable hydrogels, thus creating a platform that allows injection of sensors that do not need to be retrieved when they reach the end of their useful lifespan.

## 1. Introduction

The increasing prevalence of diabetes over the past few decades has turned this disease into one of the leading causes of death worldwide^[1]^. Afflicted individuals not only have to manage glucose levels regularly but also run the risk of long-term complications that can result in organ failure and even death^[1]^. A combination of lifestyle modifications and multiple glucose measurements throughout the day is generally used to reduce the risk of these dire complications^[1]^. Disposable glucose test strips are widely used for self-monitoring of glucose, where a droplet of blood from a finger prick is analyzed by a glucometer. However, this approach does not provide sufficient temporal information on the glucose fluctuations throughout the day, and therefore, is less effective in detecting adverse hypo- and hyperglycemic events^[2,3]^. A promising alternative to infrequent measurements is the use of continuous glucose monitoring (CGM) systems. CGM has the ability to provide a much more detailed picture of the blood glucose levels, providing critical information to the patient to be proactive in reducing dangerous spikes, which has been shown to have a positive impact on the treatment outcome and quality of life in diabetic patients^[3]^.

Although efforts have been directed towards developing non-invasive blood glucose monitoring^[4]^, most available CGMs require surgical insertion or minimally invasive procedures^[5]^. These types of sensors measure the concentration of glucose in the interstitial fluid (ISF), which has been shown to have a close correlation to blood glucose concentrations^[6,7]^. CGMs today are exclusively used by Type 1 diabetic patients but are considered too expensive, painful and difficult to manage for the far larger type 2 group of diabetic patients. Significant efforts are being undertaken to make CGM technology more available, and less complicated as well as more comfortable to use, so that this technology could eventually become the mainstream monitoring solution for diabetic patients to significantly improve patient outcomes and reduce the economic burden. Most of the CGMs in the market are based on electrochemical detection modalities. Typically, a measurable electric current is produced by an enzyme catalyzed reaction of glucose using glucose oxidase or dehydrogenase^[8],^ providing the foundation to measure glucose concentrations^[9]^. Common challenges associated with these types of sensors are the trade-off between selectivity and sensitivity, oxygen dependency, and interference of some medications and ascorbic acid (Vitamin C)^[5,10]^. More recently, development efforts are focused on boronic acid-based glucose sensing^[11]^. Glucose sensing using boronic acid does not exhibit oxygen dependency issues and can be tailored to be highly selective towards glucose^[11,12]^. Also, unlike enzymatic glucose sensing, the recognition process is based on a reversible reaction.

Among these boronic acid-containing sensors, hydrogel-based sensors are particularly attractive for *in vivo* sensing applications, in part, due to the potential biocompatibility of many hydrogel-based structures^[13]^ and the ability to tailor these hydrogels to many different analytes^[14–17]^. A body of literature demonstrated the continuous monitoring of glucose *in vivo* using hydrogel structures with boronic acid-based moieties and fluorescent monomers ^[18–22]^. In these hydrogels, the fluorescence behavior of dyes that are immobilized in the hydrogel network change when exposed to glucose. Probably the most notable case is the report from Dehennis et al.^[22]^, which eventually led to a commercial product for non-conjunctive glucose monitoring *in vivo* for three to six months^[23,24]^. However, most of the reported biosensors for *in vivo* monitoring of glucose still require implantation of electronics or other rigid materials inside the body, which may increase their complexity, reduce biocompatibility, and increase the burden and risk to the patient. The hydrogel-based approaches that require implanting electronics or rigid materials also prevent the ability to develop biodegradable hydrogel-based implants that can resorb inside the body without the need for surgical removal after its useful lifespan.

To address some of these challenges, we recently reported on a new ultrasound-based sensing platform that makes use of the resonance behavior of hydrogel structures. Hydrogels can be specifically designed to elicit a volume-phase response to changing levels of glucose, and this volume-phase transition changes the dynamic mechanical behavior of these structures, which can be probed using ultrasound waves. Hydrogels tested with this platform consisted of a hydrophilic network of acrylamide, methacrylate, and boronic acid components and have a volumetric response to glucose^[25]^. We demonstrated the working principle of this platform by measuring various concentrations of glucose as an example of an important health-related biomarker^[26]^. More importantly, this sensing principle does not require implantation of any electronics inside the body or percutaneous wire connections, and therefore, holds a significant advantage for applications in the implantable sensor field such as continuous glucose monitoring. The current work details the proof-of-concept results for monitoring glucose concentrations *in vivo* using this sensing platform and investigates its performance.

## 2. Materials and Methods

### 2.1. Fabrication of the implants

The stimuli-responsive hydrogels used in this study contain boronic acid moieties incorporated into the polymer network, enabling them to have a reversible volume response to glucose. It is shown that pH and ionic strength also create a volume response in these types of hydrogels^[26,27]^. The fabrication process for the hydrogel structures was previously described in detail^[26]^. In brief, 242.6 mg of 4-(2-hydroxyethyl)piperazine-1-ethanesulfonic acid (HEPES, Sigma-Aldrich) was dissolved in 1 L of deionized water with a pH value that was adjusted to 8.0 to make 1 mM HEPES buffer solution. Next, 19.1 mg of 3-aminophenylboronic acid (3-APB, Frontier Scientific) was dissolved in 87 µL of dimethyl sulfoxide (Sigma-Aldrich). Then, 237 µl of 30 w/w% acrylamide (Fisher Scientific) in 1 mM HEPES buffer, 193 µL of 2 w/w % N,N′-methylenebis(acrylamide) (Sigma-Aldrich) in 1 mM HEPES buffer, 20.5 µL of N-[3-(dimethylamino)propyl]methacrylamide (Sigma-Aldrich), and 309 µl of 1 mM HEPES buffer were combined and mixed by vortexing. Finally, 25.8 µL of 4 w/w% of lithium phenyl-2,4,6-trimethylbenzoylphosphinate (Sigma-Aldrich) was added to the mixture in a yellow light environment. The hydrogel structures were fabricated by exposing the prepared pre-gel solution to UV light inside their respective mold structures. The structure that was considered for this work consists of a hydrogel sheet with the thickness of 254 μm, which has pillars with the height of ~25 µm and the diameter of 60 µm spaced 10 µm from each other fixed on a 25 µm thick polyimide (PI) film, as seen in **Figure 1A**.

**Figure 1.**
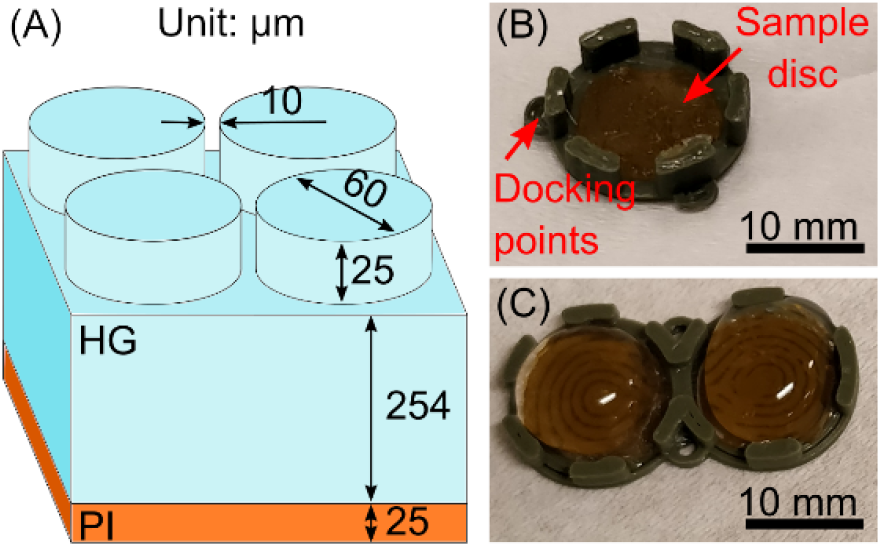
The design of the hydrogel-based structures used for implantation. (A) The dimensions of the hydrogel resonator structure that is used for this study. (B) An exemplary PLA casing with one well and one mounted sample on it (PLA-SW), (C) An exemplary PLA casing with two wells and two samples mounted in these wells (PLA-DW). Note that the hydrogel samples are under a PBS droplet to be protected from dehydration.

The prepared hydrogel structures were cut into circular discs of 8 mm in diameter using a biopsy punching tool. These discs were then fixed on a casing using a cyanoacrylate-based glue (Krazy Glue™, Elmer’s Products.). The hydrogel structures were kept hydrated throughout the preparation process and stored in a 10 mM phosphate-buffered saline (PBS) solution with a pH of 7.4 and prepared in our lab as described previously^[26]^. The casing was made of a 3D printed polyacrylic acid (PLA), which had one well (SW) or two wells (DW) to accommodate samples, as seen in Figure 1B (PLA-SW) and 1C (PLA-DW), respectively. The PLA casings had an inner diameter of 9 mm and an outer diameter of 11 mm for each well, with raised walls of 2 mm height as well as docking points around the implant. Note that the raised sidewalls surrounding the hydrogel disk are not continuous to allow for the exchange of analyte-containing biofluids from the external environment to the hydrogel structures. The total lengths of the implants are 11.0 and 20.7 mm for PLA-SW and PLA-DW casings, respectively. The PLA was chosen due to its relatively good biocompatibility for short-term implantation purposes^[28]^ and its wide availability and simplicity in prototyping. The purpose of the PLA casing was to simplify the localization of the hydrogel during *in vivo* testing and protect the resonators from possible physical impacts from its surrounding. In future work, the PLA casing might no longer be needed and be replaced with more biocompatible material options such as hydrogels with higher polymer content with embedded contrast enhancing features for the particular sensing frequency. The hydrogel resonators can also be placed inside a protective and permeable matrix or cage to avoid collapsing of tissue to compress the resonators. As it will be explained later, the PLA-SW design was only used for the two initial experiments and the subsequent experiments used a PLA-DW design to test more samples per subject.

The three most common techniques for terminal sterilization of implantable devices are autoclaving, gamma irradiation, and treatment with ethylene oxide gas^[29,30]^. All three of these techniques have been shown in many cases to affect hydrogel properties, though the precise effects must be evaluated on a case-by-case basis^[30]^. In the current study, the implants were autoclaved inside the PBS solution after preparation to ensure sterility and were stored in a sealed container at room temperature afterward. The total volume of the PBS solution for autoclaving was ~30 mL. The autoclaving process was a standard process of ramping the temperature up to 121 °C and maintaining it for at least 20 minutes at elevated pressures to avoid volume loss of the PBS solution and ensure sterilization^[31]^.

### 2.2. Experimental setup

For *in vitro* ultrasound glucose measurements, the implants were glued to the bottom of a container in a straight line and were imaged inside a PBS solution using a medical ultrasound imaging machine (ACUSON S2000, Siemens Medical Solutions USA, Inc.) Note that the container was sealed with plastic wrap during the experiments to reduce the effect of evaporation during prolonged measurement periods. Two-dimensional pulse-echo (B-Mode) images were acquired at 4 MHz on the ultrasound probe (9L4, Siemens Medical Solutions USA, Inc.) and were saved in an 8-bit grayscale format. The imaging plane for the ultrasound array was adjusted to create a cross-sectional ultrasound image of the implants similar to our previous report^[26]^. The focal length was adjusted to the minimum possible value (3 cm). The dynamic range and power were 90 dB and 100%, respectively, and all the image processing features of the equipment were turned off.

For *in vivo* glucose monitoring, healthy albino Sprague Dawley male rats (Charles River) with a weight of approximately 300 g were used as the animal model system. The subjects were anesthetized using isoflurane throughout the procedure. The abdominal area was shaved, disinfected, and locally numbed with lidocaine injections followed by a small incision exposing the subcutaneous tissue. A pocket for the placement of the sensor was created using a blunt-tip scissor. The sensor was placed in the pocket and kept in place by suturing the docking points to the skin. The implantation site was flooded with a sterile 0.9% sodium chloride solution before closing to avoid trapping air cavities underneath the skin that would interfere with the ultrasound measurements. The incision was closed by sutures and sealed using surgical glue (VetBond, 3M). Then, a water-based ultrasound transmission gel (Medline Industries) was applied to the skin and an array transducer (9L4, Siemens Medical Solutions USA, Inc.) was adjusted to obtain a cross-sectional view of the implants, as shown in **Figure 2A and 2B** using an ultrasound imaging machine (ACUSON S2000, Siemens Medical Solutions USA, Inc.) Once the best imaging position was reached, the probe was fixed in place using a laboratory stand with clamps. The imaging parameters were set identical to the *in vitro* experiments. Ultrasound images were obtained from the implant every 5 minutes throughout the duration of the experiment at 4 MHz. **Figure 2C** shows a sample image taken from a subcutaneously implanted sample, as shown in **Figure 2D**. The total time of the experiment was limited to no more than 10 h based on established Institutional Animal Care and Use Committee (IACUC) protocols. The subjects received sterile 0.9% NaCl injections subcutaneously with a dosage of 1 mL/h to maintain hydration levels. In addition, a heating mat was placed under the subjects throughout the procedure to avoid hypothermia, and the subjects were monitored for temperature using a thermocouple thermometer (Kent Scientific Corporation). The tip of the tail was pricked to obtain a small volume of blood from the capillary veins for reference glucose measurements. Two brands of commercial glucose test strips, namely Alpha Track^®^ (Zoetis Inc.) and Contour Next^®^ (Ascensia Diabetes Care), were used to measure capillary blood glucose levels approximately every 15 minutes throughout the procedure. The scab formed on the site between the measurement intervals was removed using a sterile gauze before obtaining a fresh blood sample for measurement. The blood glucose concentration was modulated during the procedure with insulin (Humulin R, Eli Lilly, and Company) and glucose (Dextrose 50% IV solution, Hospira, Inc.) injections. First, the implants were allowed to equilibrate for a minimum of 2 h. Then, the subjects received an intravenous (IV) 1-10 U/Kg insulin injection with dosage depending on the baseline glucose levels. The glucose levels and the vital signs of the subjects were closely monitored. In case of approaching dangerously low levels of glucose or the onset of seizures, which includes abnormal heart rate and shallow breathing^[32]^, IV injections of glucose were immediately administered. All procedures were performed in accordance with the approved protocol #19-01008 and under the guidance of the University of Utah IACUC.

**Figure 2.**
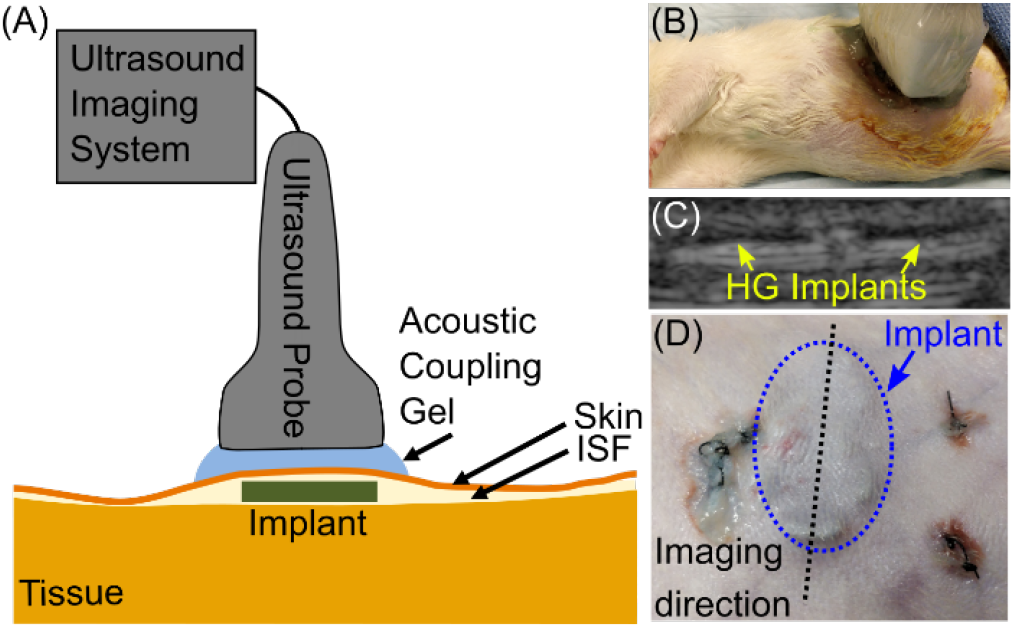
Schematic and representative images of an in vivo experiment for monitoring glucose levels in a rat model. (A) The schematic of the test setup. (B) The positioning of the ultrasound probe above the subject aligned to image the hydrogel structures implanted subcutaneously. (C) A sample B-mode ultrasound image obtained from an implant at 4 MHz. (D) An exemplary position of a sub-cutaneous PLA-DW implant.

A total of five experiments were performed in this study. The first two experiments were performed with PLA-SW. The next three experiments were performed with a PLA-DW to obtain more information using the available implantation space.

We used the abdominal region as the implantation location, which could potentially provide better glucose exchange between blood and ISF due to increased vascularity^[33]^. Experiment #5 was performed with an imaging direction depicted in **Figure 3A**, whereas the rest of the experiments were performed with an imaging direction depicted in **Figure 3B**.

**Figure 3.**
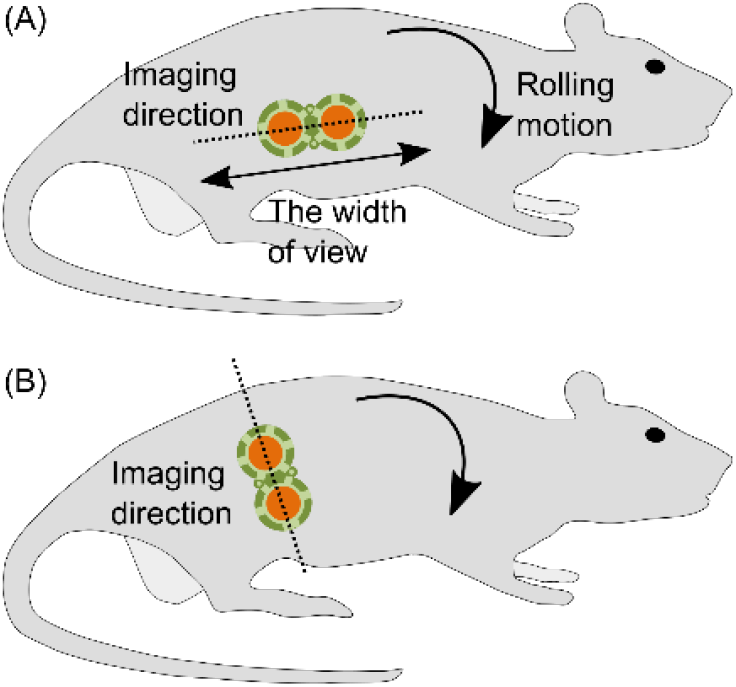
The imaging plane horizontal (A) and vertical (B) relative to the length of the body.

### 2.3. Data Analysis

The intensity of the ultrasound reflections from the hydrogel samples depends on the swelling state of the hydrogel structures and shows up in ultrasound images in the form of a local pixel intensity change in the interface of the hydrogel and the surrounding environment. Data analysis, generally in line with previous work^[26]^, was performed for both *in vitro* and *in vivo* experiments to correlate the information contained in the ultrasound images to the glucose concentration. In short, the collected images were stabilized based on an algorithm by Tevavez et al.^[34]^ that is available as an add-on script for the ImageJ^[35]^ software package. After the image stabilization, a box of a size of 100×20 pixels was selected in which the interface of the hydrogel and the surrounding environment was centered in the box. The dimensions of the box are not optimized. Doing so to contain only the pixels from the region of interest can potentially increase the signal-to-noise ratio. Nevertheless, the mean grayscale value (MGV) of the box always contains the information on the absorption of ultrasound waves by the hydrogel structure(s) as long as the position of the box is kept constant throughout all the images. With respect to the *in vivo* experiments, the disruptions resulted from the natural reflexes, movements, and relaxation of the body were observed in the MGV signal as a step change of the overall level of the signal, which was not observed in our *in vitro* experiments^[26,36]^. These disruptions were corrected using a baseline correction technique in addition to the previously applied stabilization technique. Assuming *MGV[i], MGV*_*c*_*[i]* are the signal before and after baseline correction, respectively, and *i*_*d*_ is the index of the point where the disruption happens, baseline correction involved adding the difference that the disruption has created to the baseline of all the data points after the disruption, as shown in equation 1 (see supplementary information for more details). Further, the corrected MGVs were normalized to unity. 

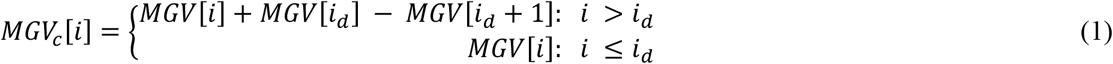

An increase in blood glucose (BG) results in a decrease in MGV, as detailed previously^[26,36]^. Therefore, after correcting for the spatial alignment for disruptions, the initial criteria for assessing the quality of the obtained MGV signal needs to include observing a negative correlation with the BG profile as established by the commercial glucose test strips. Then, the MGV signal needs to be compensated for its time lag compared to the BG profile. This time lag has resulted from the inherent delay between the blood glucose and ISF glucose levels^[37,38]^, the delay for the diffusion of the glucose from ISF to the implants, and the delay that is caused by the inherent response time of the hydrogel. Next, the MGV signal needs to be calibrated to the BG profile to create the sensor’s glucose readout in ISF (IG). These steps are explained in detail below.

The MGV signal was first interpolated linearly, and then a correlation minimization algorithm^[39]^ was employed as described below. The MGV(t) was shifted in the time domain with the parameter τ ∈ℤ, −180< τ <180 to create 361 shifted functions. Note that the unit for both τ and t is in minutes. For each shifted function, MGVτ(t), the MGV values were calculated in the temporal points corresponding to the BG signal. Note that the BG values in each time point are the mean value of the readouts resulted from the two brands of glucose test strips to increase accuracy. Calculating the correlation between the shifted MGVs and BG datasets creates a correlation function with respect to τ. The τ value that minimizes the correlation function is equal to the actual time delay between BG and MGV. After compensation for the time delay between the MGV and BG, two points in time were chosen where the BG signal was in a steady state as determined by visually inspecting the data, and the implants were given maximum possible time within the duration of the experiment to exhibit response to the second order changes in the BG profile. One of these calibration points was close to the time point where the insulin was injected. The second calibration point was closer to the end of the experiment. The BG and MGV levels were calculated for each of these settling time points using the mean value of three temporal data points in the vicinity of that time point. The two values in the MGV signal were mapped to their corresponding BG values using a linear function to create IG. The value of IG, which is a continuous function of time, was calculated in the location of matching reference BG points to create paired points of IG and BG, which was used to determine further performance metrics.

The performance of the sensors was measured by the correlation value between the IG and BG. In addition to the correlation, the Mean Absolute Relative Difference (MARD) and Clarke Error Grid Analysis (EGA)^[40]^ were used to provide a cursory evaluation of the continuous glucose sensor. The MARD value is the mean value of the absolute relative difference between the paired points from the sensing readout under investigation and a reference test^[41]^. Lower MARD values in general indicate the CGM readout is closer to the reference values. Today’s CGMs report values closer to 10^[42,43]^. Further improvement to reduce the MARD below 10 is questioned for their merit in glucose management^[3]^. The Clarke Error Grid is a chart introduced by Clarke et al.^[40]^ based on clinical data from diabetic patients, which plots the glucose readout from the sensor against reference glucose values from a proven test. The chart is divided into five sections (A-E) based on the measurement accuracy and potential risk deviating values pose to the patient’s health. Zone A represents a region where the glucose readout is considered clinically accurate, while zone B is considered to be clinically acceptable with errors that do not result in inappropriate treatment. Zones C, D, and E are indicating clinically significant errors that potentially pose a threat to the patient’s health^[40]^. Therefore, the readout from a glucose sensor needs to fall in zones A and B to be clinically accurate and safe.

## 3. Results

One of the requirements for our procedure is to use sterile implants for all *in vivo* experiments, necessitating the need to study the effect of sterilization of the hydrogel structures on the sensing performance. Three identical implants were prepared for *in vitro* studies: one with and two without autoclaving. Autoclaving was done as described in previous sections. The implants were imaged every five minutes for the duration of the experiment. The implants were exposed to different glucose concentrations, allowing six hours for each concentration step to reach equilibrium. **Figure 4** shows the *in vitro* measurements of these three samples at an ultrasound frequency of 4 MHz. The size of the selection box for calculation of mean gray value was 100×5 pixels.

**Figure 4.**
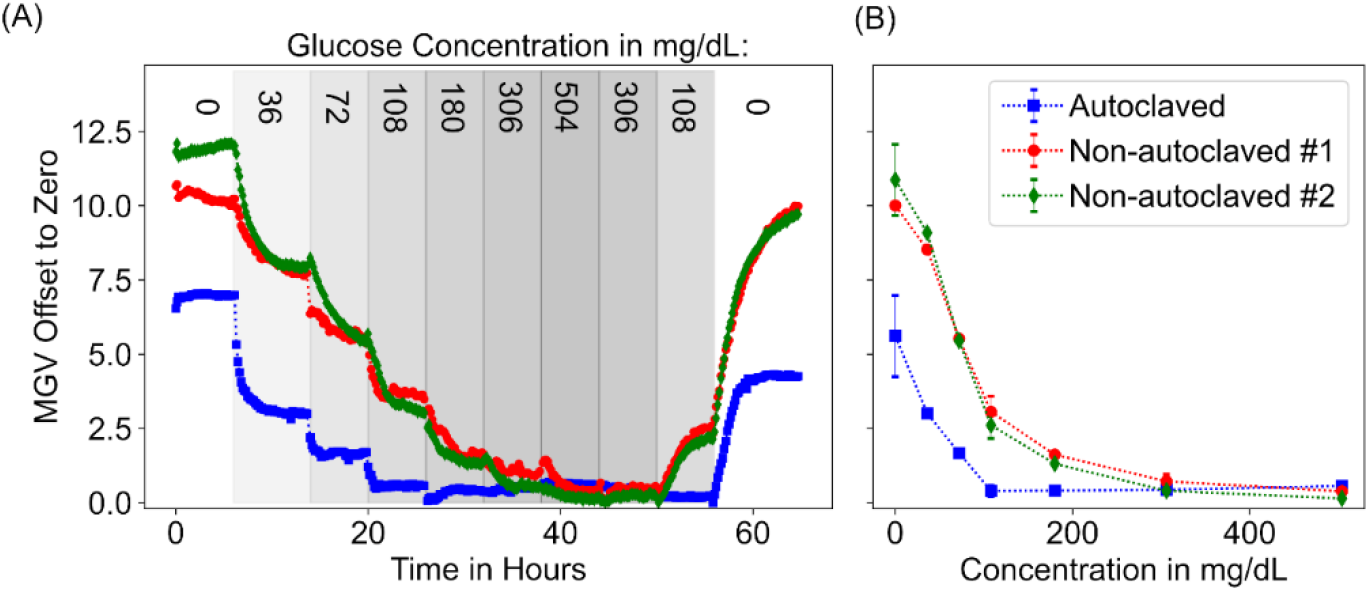
(a) Time response of samples for different concentrations of glucose at 4 MHz. (b) MGV versus glucose concentration after 6 h incubation. The error bars for the 0 mg/dL, 108 mg/dL, and 306 mg/dL concentration steps are the difference between the two data points. The rest of the points are single points. The dashed lines are guides to the eyes.

The five detailed in vivo studies (#4-8) consist of two experiments with PLA-SW (#4 and 5) and three experiments with PLA-DW (#6, 7, and 8) implants. Experiment #6 had one sample containing hydrogel pillars and one control sample consisting of an empty 25 μm PI film. Experiment #7 had one sample containing hydrogel pillars and one sample consisting of a flat 254 μm hydrogel sheet on a 25 µm PI film to further verify the unique response of the hydrogel pillar samples at 4 MHz. Overall in these experiments, the MGV signal was obtained from six instances of hydrogel pillar structures (#4, 5, 6.2, 7.2, 8.1, and 8.2) and two instances of other samples (#6.1 and 7.1). The obtained images from these experiments were corrected for their baseline and then were normalized to unity. Figure 5 shows the resulting MGV signals for all the instances (see supplementary information for the MGV signals before applying the corrections). Next, the MGV signals were corrected for time lag and then were calibrated. The metrics that were used for evaluating the functionality of the implants were the time delay between BG and IG, correlation of IG with BG after compensation for time lag, MARD value, and the clinical accuracy of the paired points in the EGA plot, as shown in Table 1.

**Table 1.** Summary of correlation, time delay, MARD and standard deviation of MARD, the number of paired points, and EGA for each case. The column for the EGA Accuracy lists three numbers for each experiment representing the percentage of total data points in zone A and B combined and in parenthesis the individual contribution of each zone, respectively. Grayed rows show the control samples.

| Experiment | Casing Type | Correlation (%) | Delay (min) | MARD ± 95% CI | Number of Paired points | EGA Accuracy A, B Zones (%) |
| --- | --- | --- | --- | --- | --- | --- |
| 4 | PLA-SW | -85 | 53 | 31 ± 5 | 38 | 95 (71, 24) |
| 5 | PLA-SW<sup>a)</sup> | 21 | 150 | 165 ± 58 | 23 | 36 (18, 18) |
| 6.1 | PLA-DW | 37 | 21 | 167 ± 114 | 59 | 57 (29, 27) |
| 6.2 | PLA-DW | -91 | 33 | 43 ± 7 | 57 | 82 (64, 18) |
| 7.1 | PLA-DW | -38 | 2 | 308 ± 226 | 50 | 56 (34, 22) |
| 7.2 | PLA-DW | -91 | 51 | 42 ± 6 | 41 | 83 (68, 15) |
| 8.1 | PLA-DW | -87 | 134 | 62 ± 8 | 42 | 88 (73, 15) |
| 8.2 | PLA-DW | -91 | 118 | 52 ± 8 | 44 | 96 (67, 29) |

a) Imaging was performed horizontal relative to the length of the subject's body, which resulted in a signal loss.

**Figure 5.**
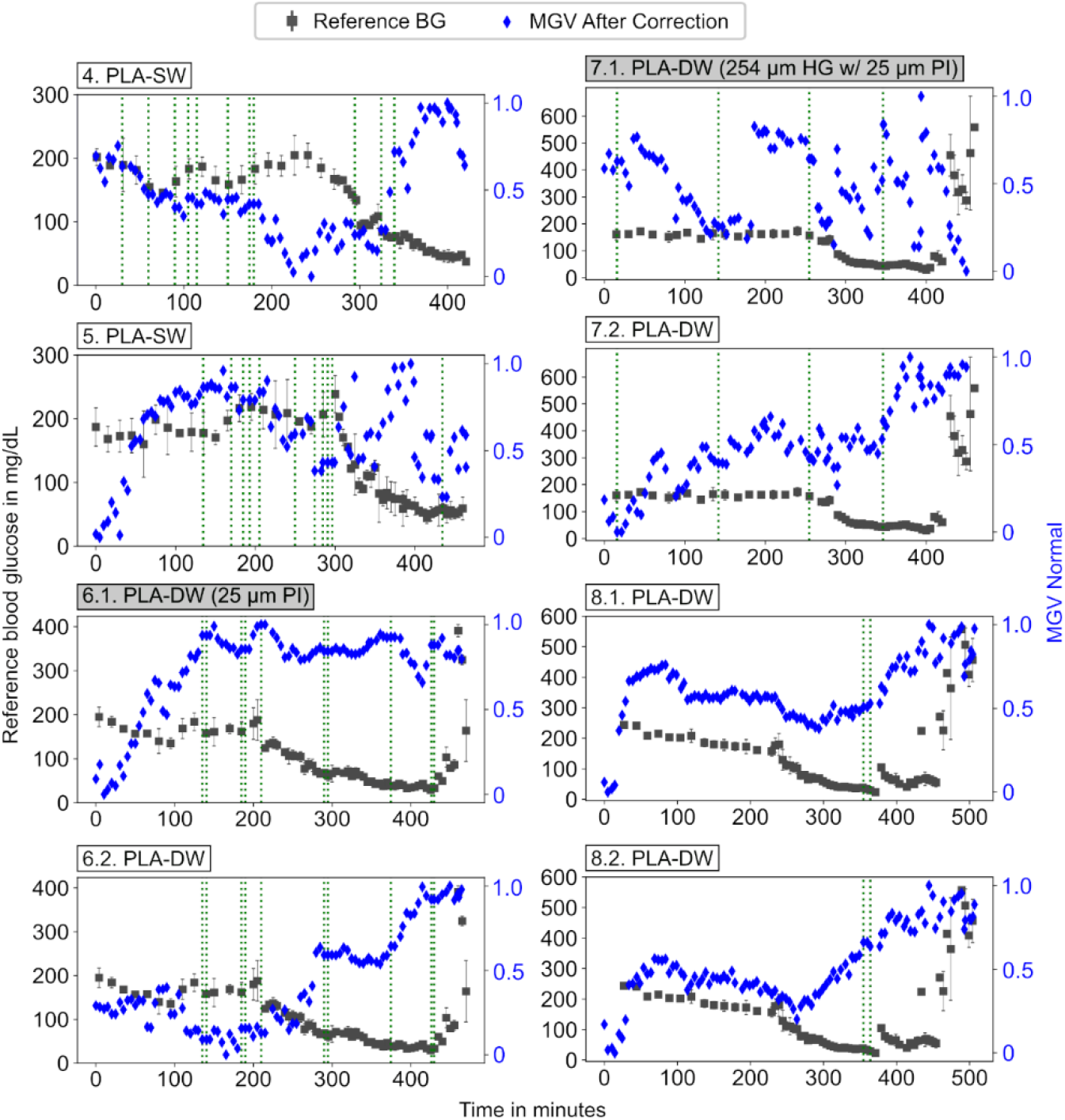
MGV signal after correction versus the reference BG profile for the eight instances of the samples tested in the subjects. The vertical green dashed lines indicate time points where corrections for probe disruptions were applied. The error bars for the BG profile are the difference between the two measurements made with two different brands of blood glucose meters. The grayed headings show instances that are not hydrogel resonator samples.

Except for experiment 5, which had a signal loss due to its imaging direction and the control samples, the rest of the calibrated IG signals and their corresponding EGA plots are shown in **Figure 6**.

**Figure 6.**
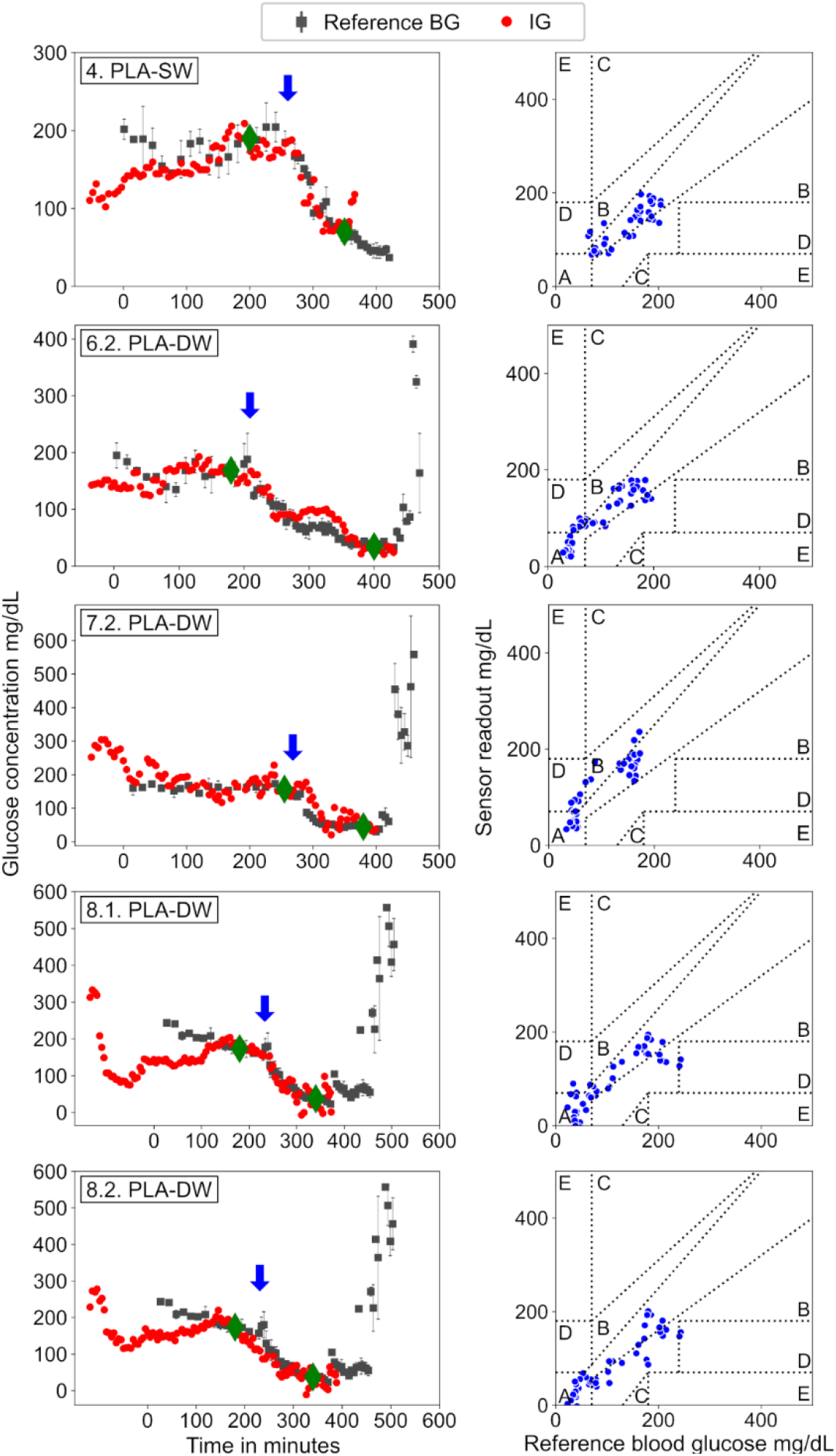
The temporal plots for comparison of IG measured using the implanted sensors with the reference BG values measured by glucose test strips and their corresponding Clarke Error-Grids. The error bars for the reference signal are the difference between the two measurements made with two brands of blood glucose meters. The blue arrows show the approximate location of the insulin IV injection. The green diamonds indicate the location of the calibration points.

## 4. Discussion

The results presented in Figure 4 show that samples that were autoclaved and that were not autoclaved follow a similar trend, which indicates the smart hydrogels remain responsive to glucose after autoclaving. However, autoclaving does result in a decrease in the ultrasound signal intensity at a fixed frequency of 4 MHz. According to a review by Galante et al.^[30]^, numerous studies have shown that autoclaving of synthetic hydrogels often changes their properties, in particular the elastic shear modulus G. Given that the hydrogel swelling ratio and the resonance frequency of the smart hydrogel resonators both depend on the value of G, the reduction in the sensor signal due to autoclaving could be attributed to a shift in resonance response of the smart hydrogel structures and a reduction in their swelling response. The response time of the hydrogel structures was also compared by calculating their T_90_ value using an exponential fit^[36]^ when transitioning from an external environment containing no glucose to 2 mM glucose. The transition to higher glucose levels results in a negative correlation with MGV, as reported before^[26]^. The T_90_ response time was 2.2 h (R^2^=98.8%) for autoclaved, and 5.1 h (R^2^=98.9%) and 3.5 h (R^2^=99.9%) for non-autoclaved samples. Drawing a conclusion on response time based on limited samples might be prone to error; however, the data suggests that the autoclaving has caused an improvement in response time in these hydrogel structures. The reason for this possible effect is unknown and merits further investigations.

Please note that there was an air bubble trapped between the ultrasound probe and the samples in the beginning of the experiment, which was removed in the next step when transitioning from 0 to 36 mg/dL. The presence of this bubble caused a shift in the first step’s baseline. This distortion was corrected for by adjusting the baseline for the first step so that the last point in the first step matches the first point in the next step. An automated flow delivery setup could possibly reduce the occurrence of these types of experimental artifacts.

During the early experiments, we observed that the rolling motion in the subject’s body is the most common form of body movement in the subjects. An imaging direction vertical to the subject’s body, as shown in Figure 3, has a better chance to capture these disruptions. In contrast, when the imaging direction is horizontal to the subject’s body, these disruptions become challenging to detect, and also they can move the implant out of the field of view of the probe entirely leading to signal loss. As shown in Figure 5, all the experiments with hydrogel resonator samples seem to show a negative correlation to the BG profile except experiment #5. Experiment #5 was performed with an imaging direction that was horizontal to the length of the body, in contrast to the rest of the experiments, which were performed with a vertical imaging direction. During experiment #5, there were numerous disruptions to the probe, which eventually obscured the signal. This could explain the poor correlation between BG and MGV in this experiment. In all the other experiments with hydrogel resonator structures, the MGV signal has a negative correlation with the BG profile. In the case of the 25 µm PI sample (experiment #6.1) and 254 µm hydrogel with 25 µm PI sample (experiment #7.1), the MGV signal does not seem to have correlation with the BG profile. The fact that none of the samples that are not hydrogel resonators structures do not show a meaningful response to glucose supports that the response obtained from the hydrogel resonator samples are specific to these structures. All the hydrogel resonator structures except experiment #5, showed MARD values of ~ 30-60 and accuracy of ≥83% in EGA, as seen in Table 1. Although comparing the MARD value of different CGMs reported in literature is not possible due to their different testing conditions, the reported values usually range from ~10-24 with the hypoglycemic region being the least accurate^[42,43]^. Note that in the EGA analysis, in addition to the overall percentage of points in zones A and B, it is possible to divide the reported percentages for zones A and B into three regions: <70 mg/dL, 70 to 180 mg/dL, and >180 mg/dL^[44]^. However, since the number of data points in our case is limited compared to more long-term studies, we report the overall percentage. In these experiments as shown in Figure 6, the matched points that are not in region A or B, fall in region D. The majority of these points fall in the upper D region, in which the readout is over-predicting the blood glucose concentration. This could possibly be resulted from the higher order nature of the calibration curve as seen in Figure 4. As explained later, we chose first-order polynomial for calibration of sensors. Using first order polynomial for calibration when the actual calibration curve is from higher order can cause over prediction of the glucose values around 70 mg/dL and higher pushing the points to the upper D region. This behavior is more prominent in the temporal signal from the experiment # 6.2 and 7.2 depicted in Figure 6. The MGV signal and the metrics resulting from the control samples indicated that the control samples do not show a meaningful response when compared to the reference glucose measurements. This result is in agreement with the previous studies^[26]^ and supports our sensing principle.

Some considerations need to be taken into account for the MARD values presented in Table 1. First, long-term studies^[43]^ have shown that the MARD values improve weeks after implantation, possibly due to initial biofouling from the implantation procedure, which requires time to heal. Biofouling is expected to contribute to a higher MARD value in our short-term experiments. Second, an accurate MARD value estimation depends on the accuracy of the calibration curve. A variety of calibration algorithms can be used for implantable glucose monitors^[45]^. However, due to the limited duration of our experiments (≤ 10 h), which provided a challenge for obtaining more than two settling points in the data, and for simplifying the approach, we used a first-order polynomial fitting function alongside two settling points for the calibration of the implant’s readout. In future studies, longer durations allow for using more settling points to help increase the estimation accuracy of the calibration curve, which should lead to improved MARD values. Third, this study was potentially also impacted by a reference error as the point-of-care glucose test strips used have an inherent error^[46,47]^. In addition, the procedure used to extract blood samples can introduce measurement errors^[48]^. While the glucose test strips used in this study are an acceptable reference in early proof-of-concept studies, a more clinically accurate diagnostic approach^[49]^ is needed to assess the performance of the implants in future studies. Fifth, the study design including the number of data points, the number of glucose swings in the data as well as the amount of time that the sensor remains in hyper- or hypoglycemic conditions are important factors affecting the accuracy of the MARD value^[41,50]^. Here, the time allowed for each animal under anesthesia was restricted, which limited the number of collectable data from each subject. Longer experiments with more paired points well-spread in all the glycemic regions can give a better picture of the MARD values for our sensors. Lastly, the hydrogel structures reach a saturation response after 6 mM (128 mg/dL) of glucose in an *in vitro* setting, which is lower than the glucose concentrations experienced in the rat model. Although it is difficult to predict the impact of this on the glucose response in an *in vivo* setting due to a vastly more complex environment, this could possibly negatively impact the MARD value in an experiment where the subject is exposed to a prolonged hyperglycemic state.

The time delay between IG and BG signal is associated with three factors: the delay in the transport of glucose between the vascular system and the ISF, which is about 5-15 minutes^[37,38]^, the delay for the diffusion of glucose from ISF to the implant area, and the diffusion of the glucose inside the hydrogel network. The diffusion of the ISF glucose inside the sensor casing could be improved by optimizing the geometry of the implant casing and the hydrogel samples to minimize this space, and therefore, reduce the diffusion distance for the glucose molecules to reach hydrogel structures placed inside the casing. The last source of delay is the inherent delay in hydrogel response itself. Although the response times for the hydrogel samples used in this study are much higher than the biologically relevant delays, a sudden step increase in blood glucose does not typically happen in an *in vivo* setting. These continuous rather than discrete changes in the blood glucose help the sensor to better track glucose variations *in vivo* compared to the step response *in vitro*. In addition, the most significant portion of the volume response in the smart hydrogel occurs in the very early stages due to their decaying exponential function response^[51]^. In this respect, the sensor quickly reacts to the changes in glucose concentration in its surroundings and follows the trend of the always-changing glucose levels *in vivo*. Adequate computational modeling techniques can be used to retroactively predict the BG from the sensor readout for dosing decisions for closed-loop insulin delivery systems^[52]^. The wide variation in time delays in different experiments and the fact that all the hydrogel samples have relatively identical response times indicate that factors outside the hydrogel response itself contribute to the variation in sensor performance. These factors could include possible physiological variations between the subjects, the location of the implant, access to ISF, and the size of the pocket that was created for the subcutaneous placement of the implant.

Considering that the current study is only a proof-of-concept demonstration and the limited data sets used in the evaluation, we cannot draw definitive conclusions on the performance of these sensors compared to other reports. However, the available data suggests that this platform has the potential to be a great addition to the continuous glucose monitoring field.

## 5. Conclusion

In this paper, we demonstrated the feasibility of remote glucose monitoring *in vivo* using ultrasound waves to query subcutaneous implants based on glucose-sensitive smart hydrogels resonator structures. We first presented the fabrication and assembly process for the implants and then validated the glucose response of the smart hydrogel structures after sterilization through autoclaving. The fabricated implants that only consisted of the plastic casing and smart hydrogels were then implanted *in vivo* and imaged remotely using a conventional ultrasound imaging system to track the glucose levels in the implantation site for a maximum of 10 hours. Although longer *in vivo* studies will provide a better performance assessment, using common performance metrics for glucose sensing that included sensing delay, correlation, MARD, PARD, and EGA, we have shown that glucose sensing that is based on resonant absorption of ultrasound waves is a promising approach for *in vivo* sensing. Further optimization of these hydrogel sensors for faster response (structural optimization) and increased glucose sensitivity over a larger dynamic range (chemical optimization) will pave the way to significantly enhance sensing performance characteristics.

## Acknowledgment

The authors would like to thank Nassir Marrouche, David Warren, and Stephen Wasmund for helpful discussions as well as Denis Parker for providing access to the ultrasound imaging system. The authors also thank the members of the Institutional Animal Care and Use Committees committee and clinical veterinarians for providing guidance on best practices for animal studies and Hunter Strathman for helping with the initial training.

Funding from the Joe W. and Dorothy Dorsett Brown Foundation, the Olive Tupper Foundation, and Sentiomed, Inc. is gratefully acknowledged.

This work was performed in part at the Utah Nanofab, which is sponsored by the College of Engineering, Office of the Vice President for Research, and the Utah Science Technology and Research (USTAR) initiative of the State of Utah. The author(s) appreciate the support of the staff and facilities that made this work possible.

