## Supporting Information for "*In vivo* Monitoring of Glucose using Ultrasound-induced Resonance in Implantable Smart Hydrogel Microstructures" (from another version of this preprint: version 1, 2021-04-11)

**Content:**

- Criteria for detection of disruptions to the probe relative to the samples
- Baseline adjustments for mean grayscale value signals from the samples
- Example data analysis for calculating the correlation and time lag

### Criteria for detection of disruptions to the probe relative to the samples

During the *in vivo* experiments, controlling the natural reflexes, movements, and relaxation of the body is difficult. Therefore, disruptions to the relative location of the probe and the implant is inevitable. Many of roation and translation movements (x-y movements) can be corrected with image stabilization techniques. However, some disruptions cause a shift in the signal baseline, which is usually caused by a disruption in the z-axis. Therefore, the signals need to go through a signal-leveling step to compensate for disruptions and signal shifts caused by body movement, before further processing. This step involves shifting the signal baseline before and after each disruption point to match the two points in question to re-establish baseline continuity as described mathematically in the main text. Figure S1a depicts the process applied to a set of hypothetical data points. The time point at which a considerable disruption occurs to the probe location relative to the implant was chosen manually. First, a list of suspected time points for disruptions was chosen by observing every image manually and comparing them with their previous image to find abrupt noticeable changes. Then, the change in the amplitude of MGV in the raw signal obtained from the implants was examined at the suspected time points. If an anticipated disruption was creating a step change in the signal, the points before and after disruption were corrected by aligning their baseline signal. However, if the time points in question did not create a step response in the signal level, the point was not considered a disruption. Note that this procedure was used only in extreme cases to preserve the integrity of the overall signal. Figure S1b shows two examples of consecutive ultrasound images, which highlight a disruption of the probe location that created a baseline change in the signal and needed to be corrected. In these images, the implant's cross-sectional width and its representation changed abruptly. These changes indicate disruptions to the probe that are in z direction (off-plane). These disruptions are impossible to correct for using the rigid body transformations which can only correct the in-plane (x and y) movements.

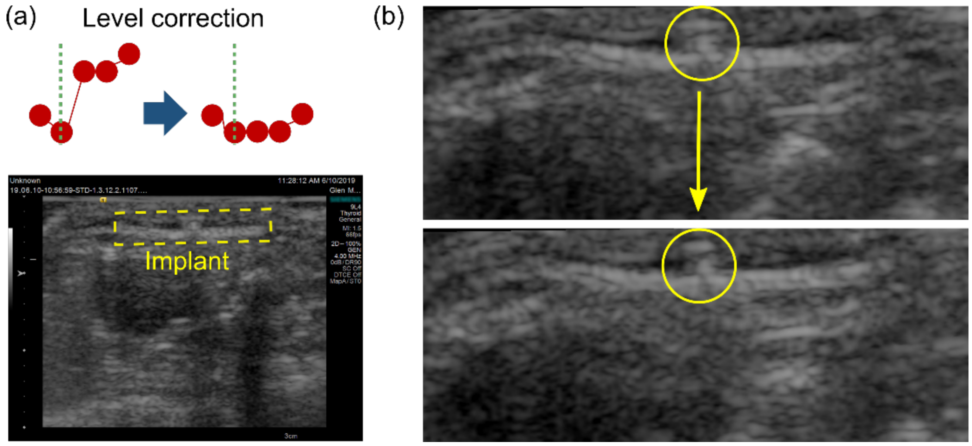


Figure S1. (a) representation of a bassline shift on hypothetical data points. (b) A sample instance that disruption created a bassline shift in the signal and needed to be corrected. As seen in the circled area, the intensity of a feature corresponding to the implant has dropped in the consecutive image.

In the future, automated image analysis software can be developed to localize and track the implant's location and apply baseline corrections as it detects disruptions to the probe location relative to the implant. In addition, having a dedicated reference indicator on the implant and establishing a differential measurement scheme can help remove many of these effects. Lastly, a portable and miniature ultrasound readout module fixed on the skin above the implant can significantly reduce the impact of body movements and reflexes on the obtained sensing signal, since this scheme will largely prevent the implant and the transducer from moving relative to each other due to body movements.

**Baseline adjustments for mean grayscale value signals from the samples**

Out of the eight subjects tested for this study, the first three were used for initial experiment design to evaluate experimental setup, determine optimal procedures, and to decide on the the best parameters for the study. The signal obtained from the subsequent five experiments went through a baseline correction step. Figure S2 shows the MGV signal before and after the baseline correction for each subject.

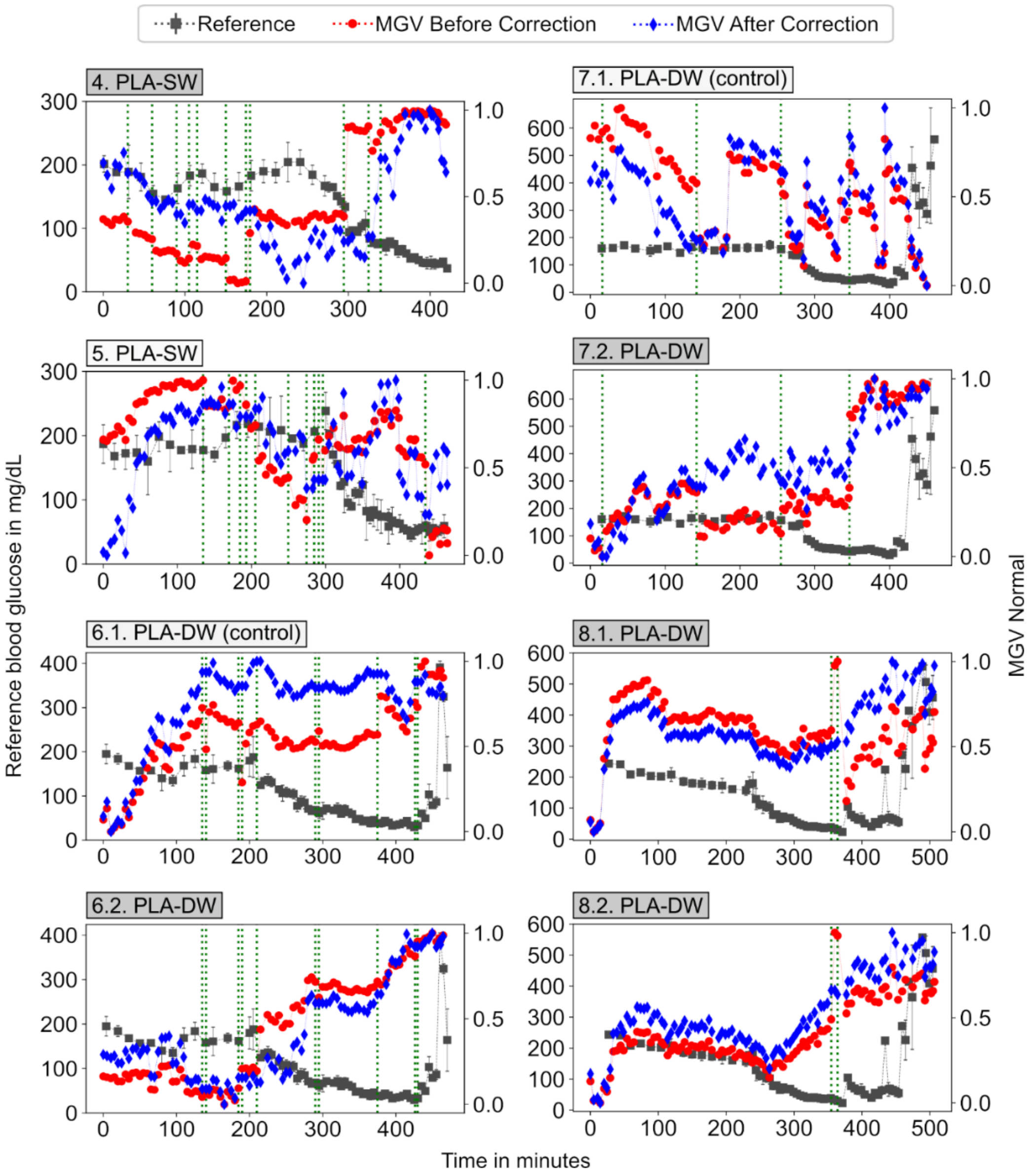


Figure S2. The normalized MGV signal before and after baseline corrections for experiments # 4 to 8. The green dashed vertical lines indicate the location of identified disruptions and subsequent signal corrections.

### Example data analysis for calculating the correlation and time lag

The points in mean grayscale value (MGV) and the points in reference blood glucose (BG) do not occur at the same time-points due to an inherent delay in glucose levels beteen blood and interstitial fluid and sensor response. In order to calculate many of the performance metrics, the value of MGV needs to be measured at the exact time when BG measurments happen. Therefore, the MGV signals that are corrected for disruptions were first linearly interpolated to create a continuous signal. This interpolation consisted of straight line segements connecting every two adjacent points. The value of the signal was then calculated at the time points corresponding to the reference blood glucose levels. Using the obtained set of paired points, we calculated the correlation between the pairs. The same procedure was carried out on the data sets when the MGV signal was shifted in time relative to the reference signal by 1-minute steps for a total of 300 minutes in both directions. Since the MGV signal is expected to negatively correlate with the reference blood glucose profile[1,2], the time shift during which the minimum occurs represents the estimated time lag between the reference blood glucose profile and the MGV signal. The minimum of the correlation graph was calculated by fitting the graph first to a 100-order polynomial function to find an approximate location of the minimum and then fitting a 20-order polynomial around the location of the minimum to find the exact point where the minimum happens. The fits were inspected visually to ensure their quality. The Figure S3 shows a correlation plot for the data set from experiment #7 and the corresponding minimum value calculated by two steps of polynomial fits. The correlation value at the minimum point is the reported correlation value between the MGV and BG. The shifting time point where this minimum happens is the time lag between MGV and BG. The MGV signal was then shifted backwards relative to the BG based on the calculated time between MGV and BG.

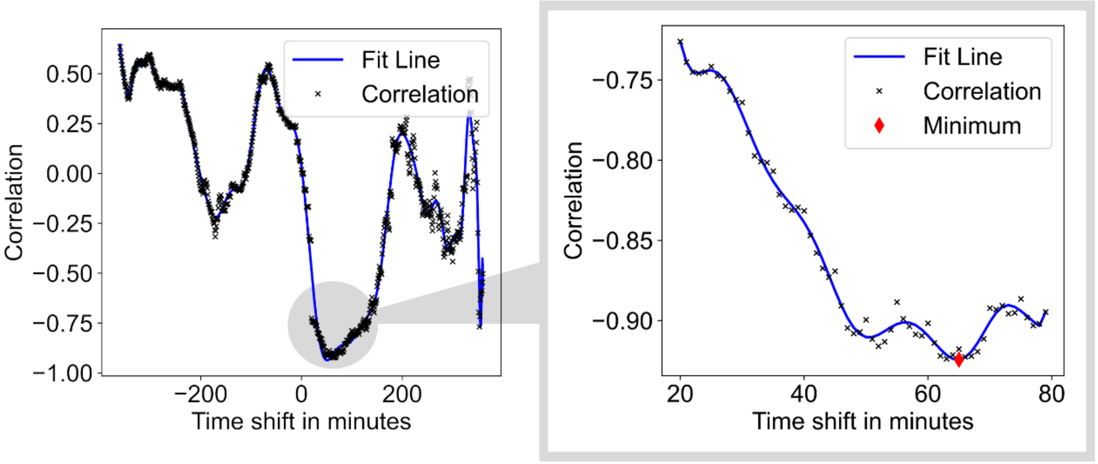


Figure S3. Reference blood glucose and MGV signal's correlation plot versus time shift. Inset on the right shows the 20-order polynomial fit for 60 points around the local extremum found in the previous 100-order fitting.
